# Serpentine mountain uplift in northern Japan triggered the divergence of a narrow endemic from a widespread (sub)arctic Asia-Alaska species complex of *Lagotis* (Plantaginaceae)

**DOI:** 10.1101/857441

**Authors:** Atsushi Sugano, Tomoko Fukuda, Yoshinori Murai, Olga A Chernyagina, Suyama Yoshihisa, Yoshihiro Tsunamoto, Hayato Tsuboi, Yoko Nishikawa, Takashi Shimamura, Hiroko Fujita, Koh Nakamura

## Abstract

In the circumboreal region, plants often have extremely-wide species ranges. *Lagotis minor-glauca* species complex widespread from (sub)arctic Asia to Alaska, however, have two allied narrow endemics in northern Japan: a serpentine plant *L. takedana* endemic to the Yubari Mountains (Mt. Yubari) and a non-serpentine plant *L. yesoensis* endemic to the Taisetsu Mountains (Mt. Taisetsu). Elucidating their origins sheds light on drivers for secondary-speciation of widespread circumboreal plants. To infer phylogenetic distinctiveness of two narrow endemics with those related taxa, which contained 25 out of all the 29 species of the genus, chloroplast DNA (cpDNA), nuclear ribosomal (nrITS), two low copy nuclear gene (LCN) markers and genome-wide single-nucleotide polymorphism genotyping (MIG-seq) were used. In the result of cpDNA analyses, the *Lagotis minor-glauca* species complex formed a clade. Within the clade, *L. yesoensis* and a portion of *L. glauca* samples formed a subclade. However, monophyly of each of the four species was not supported. In the results of nrITS and two LCN analyses, *L. takedana* was monophyletic, while monophyly was not recovered for each *L. yesoensis, L. glauca*, and *L. minor*. Based on a Bayesian dating analysis using nrITS data, the age of the most recent common ancestor of *L. takedana* was Ma (95% confidence interval: 0.05-1.75 Ma). Possible scenario is that an ancestral linage being adapted to serpentine soils migrated into the alpine habitat of Mt. Yubari, that was formed with mountain uplift by the early Pleistocene, and subsequently reproductively isolated from non-serpentine populations and speciated. The contrasting result of *L. yesoensis*, that was phylogenetically indistinct, is possibly explained by incorrect taxonomy, or alternatively, shallow history and incomplete lineage sorting. In Mt. Taisetsu, massive volcanic eruptions had occurred the Early Pleistocene and even after the last glacial period, suggesting that alpine plants have not migrated into and established populations in Mt. Taisetsu until very recently. To fully resolve the phylogeny of the three species *L. yesoensis, L. glauca*, and *L. minor*, further analyses using high resolution molecular markers are needed. The present study illustrated that two narrow endemics in northern Japan diverged from the widespread species include phylogenetically distinctive and indistinctive species, owing to historical orogeny and ecological factors.

## Introduction

In the circumboreal region, plants often have extremely-wide species ranges. These plants are adapted to harsh but less competitive environmental having one of the most severe climates on Earth where plant can survive. Despite their extreme environmental harshness, the plant species diversity is minimum in the northern areas and decreases from the equator towards the poles (Schluter and Ricklefs 1993; Huston 1994; Rosenzweig 1995; Qian et al. 1998; Qian 1999), which may be due to less ecological variation in the periphery of the circumboreal region.

Some widespread species in the circumboreal region, however, have closely related species distributed extremely locally restricted ones (narrow endemics). Geographic range sizes can vary enormously even between widespread species and closely related species (Brown et al. 1996; Gaston and Chown 1999; Webb and Gaston 2003). Such interspecific variation in range size can be associated with geographical isolation, and may in part be due to the biological traits and ecological requirements of particular species (Cowling et al. 1994; Desmet and Cowling 1999; Médail and Verlaque 1997; Hedge and Ellstrand 1999; Lavergne et al. 2003; 2004; 2005). Since geographic isolation might provide barrier to gene flow, given that marginal populations of widespread species are more possible to be isolated one, they can play an important trigger in speciation and those might be a common way by which narrow endemics originate.

This process of plants in the circumboreal region is often induced by range shift in the Pleistocene. Since the origin of species in circumboreal regions during the climate cooling in the Pliocene (Murray 1995), their current distribution have been formed by expanded and retreated of habitat and population as a result of the cycle of glacier and inter-glacier periods of the Pleistocene (Abbott et al. 2000; Avise 2000; Hewitt 2000; Abbot 2008). The range contractions of widespread species have left isolated populations in marginal areas, leading to speciation of narrow endemics. Recent molecular studies have revealed that not only climatic change in the Pleistocene, but also historical orogeny have played an important role in major trigger of species diversity (e.g. Richardson et al. 2001; Hoorn et al. 2010; Favre et al. 2014; Liu et al. 2014; Wen et al. 2014); some plants in the circumboreal region, also distributed as well as high mountains in Northern Hemisphere (Hultén and Fries 1986). Climatic changes associated with historical orogeny can be served as fragmentation of the distribution ranges of species, which might lead to barrier to gene flow (Ren et al. 2016). This process can lead to allopatric divergence and ultimately speciation (Mayr 1963; Rice and Hostert 1993).

Ecological factors, such as divergent habitats, also play significant roles in speciation (Rundle and Nosil 2005). Adaptation to newly emerged niches can stimulate populations to diverge and become isolated; ecological speciation (Filatov et al. 2016). Therefore, speciation process cannot simply be recognized as climatic changes and historical orogeny, but also ecological factors (Rieseberg and Willis 2007). Evaluating the relative roles of climatic changes, historical orogeny, and ecological factors in plant diversification lead to a better understanding of evolutionary history of narrow endemics that sheds light on drivers for speciation of species.

Since, East Asia was not heavily glaciated during the Pleistocene (Frenzel et al. 1992), many arctic and alpine plants have been expanded their range in this region. The high mountains in central Japan represents the southernmost range limit of those. Specifically, one-third of the Japanese alpine plants are endemic to Japan with closely related arctic species, or have extremely wide distribution range in the Arctic (Shimizu 1982; 1983). In this study, we explored this issue in two Japanese alpine narrow endemics of *Lagotis takedana* Miyabe et Tatew. and *L. yesoensis* (Miyabe et Tatew.) Tatew. (Plantaginaceae). *Lagotis takedana* is a serpentine narrow endemics at Mt. Yubari, Hokkaido, Japan. *Lagotis yesoensis* is also a narrow endemic at Mt. Taisetsu, Hokkaido, Japan (Fig. 1). These narrow endemic species is distributed in northern Japan that located around the distribution range of two closely related widespread species, *L. glauca* Gaertn. and *L. minor* (Willd.) Standal.. *Lagotis glauca* Gaertn. found in Rebun Island and the mountain ranges of central Honshu (Yamazaki 1993), and also in the Kuriles, Kamchatka, the Aleutian, and Alaska (Takahashi 2015; Fig. 1). *Lagotis minor* is distributed from northeastern Russia to Alaska and the Yukon (Fig. 1). Key morphological characters separating these four species are summarized (Table 1): corolla color, lower lip shape/size, stamen-adnate position, anther color/size, filament length, and leaf shape. Taxonomic treatment of the two narrow endemics as independent species is basically accepted in Japan (Yamazaki 1981, 1993; Yonekura and Kajita 2003-; Ohashi 2017); however, several studies have proposed different taxonomic treatments (Table 2). *Lagotis takedana* is sometimes treated as an intraspecific taxon of *L. glauca*: subspecies *takedana* (Miyabe et Tatewaki) et Nosaka ex Toyokuni (Toyokuni 1960; Nosaka 1974; Toyokuni 1977) or variety *takedana* (Miyabe et Tatew.) Kitamu. (Kitamura and Murata 1957; Ohwi 1965, 1978). *Lagotis yesoensis* is sometimes treated as an intraspecific taxon of *L. stelleri* Turcz., a synonym of *L. minor*: variety *yesoensis* Miyabe et Tatew. (Miyabe and Tatewaki 1933; Ohwi 1965, 1978) or subspecies *yesoensis* (Toyokuni 1977), or alternatively treated as an intraspecific taxon of *L. minor*: variety *yesoensis* (Miyabe et Tatewaki) Toyok. (Toyokuni 1981). Furthermore, in the Plant List (2013) *L. takedana* and *L. yesoensis* are treated as “unresolved” species, and *L. glauca* is synonymized with *L. minor* despite the morphological differences (Vikulova 1995; Table 1). Morphology-based taxonomy of these species is not settled yet. Molecular phylogenetic analyses of *Lagotis* are limited, and of the above-mentioned species only *L. glauca* (from Commander Islands) and *L. minor* (from Alaska) have been studied and formed a clade based on combined nrITS and cpDNA data (Li et al. 2014).

**Table 1.** Distribution area and key morphological characters of three *Lagotis* species in Japan and allied *L. minor* (Yamazaki 1981, 1993; Vikulova 1995; Ohashi 2017)

| Species | Distribution | Corolla color | Lower lip shape/size | Stamen adnate position | Anther color/size | Filament length | Leaf shape |
| --- | --- | --- | --- | --- | --- | --- | --- |
| *L. takedana* | Mt. Yubari, Hokkaido | White or pale lilac | Lower lip lobes lanceolate, as long as upper lip | Adnate to base of upper corolla lip, much shorter than upper lip | Paler color, ca. 1.4 x 1.8 mm (L x W) | Ca. 0.5 mm | Widely ovate or elliptic |
| *L. yesoensis* | Mt. Taisetsu, Hokkaido | Bluish purple | Lower lip lobes narrowly elliptic and rounded, shorter than upper lip | Adnate to middle of upper corolla lip, similar length to upper lipcorolla | Blackish blue, ca. 1.5 x 1.7 mm | Ca. 1 mm | Ovate, elliptic or oblong |
| *L. glauca* | The northern Pacific rim. In Japan, Rebun Island and the mountain ranges of central Honshu | Bluish purple | Lower lip lobes narrowly oblong and rounded, shorter than upper lip | Adnate to base of upper corolla lip, much shorter than upper lip | Dark purple, ca. 1.0 x 1.5 mm | Ca. 0.5 mm | Orbicular-ovate to widely ovate |
| *L. minor* | Northeastern Russia to Alaska and the Yukon | Bluish or whitish | Lower lip lobes acute or obtuse, slightly shorter than upper lip | Exceeding upper lip | Blue, 1-1.6 mm wide | Ca. 2-4 mm | Lanceolate to elliptic |

**Table 2.**
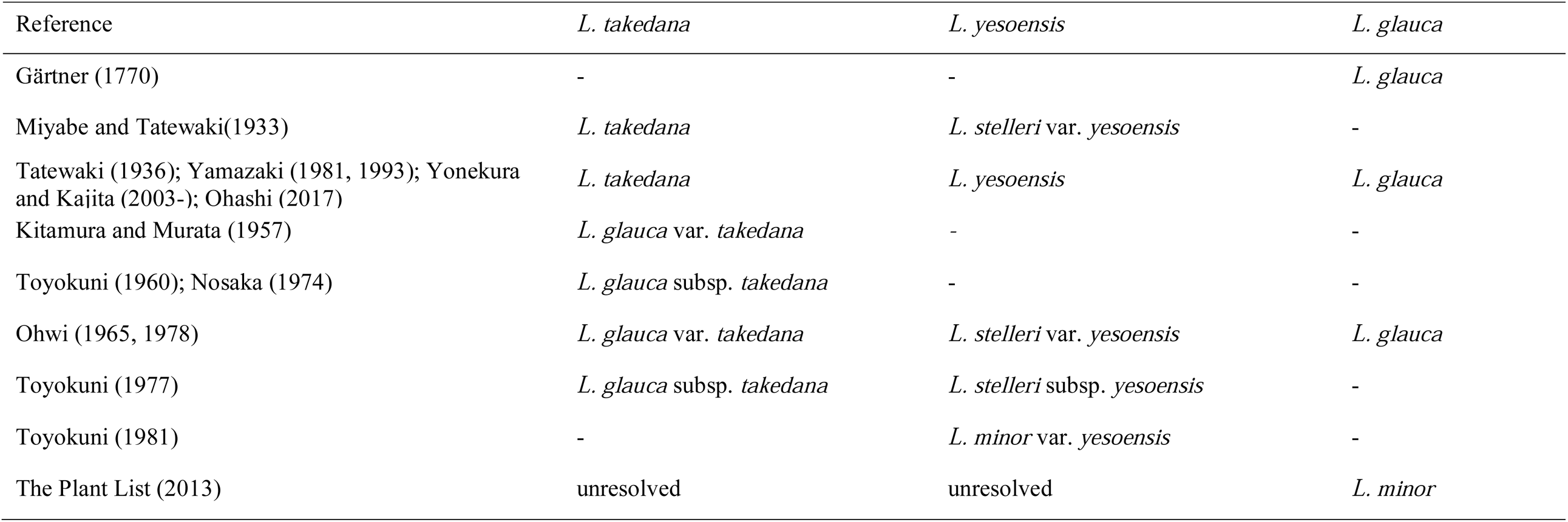
Different taxonomic treatments of *L. takedana, L. yesoensis*, and *L. glauca*.

**Fig. 1.**
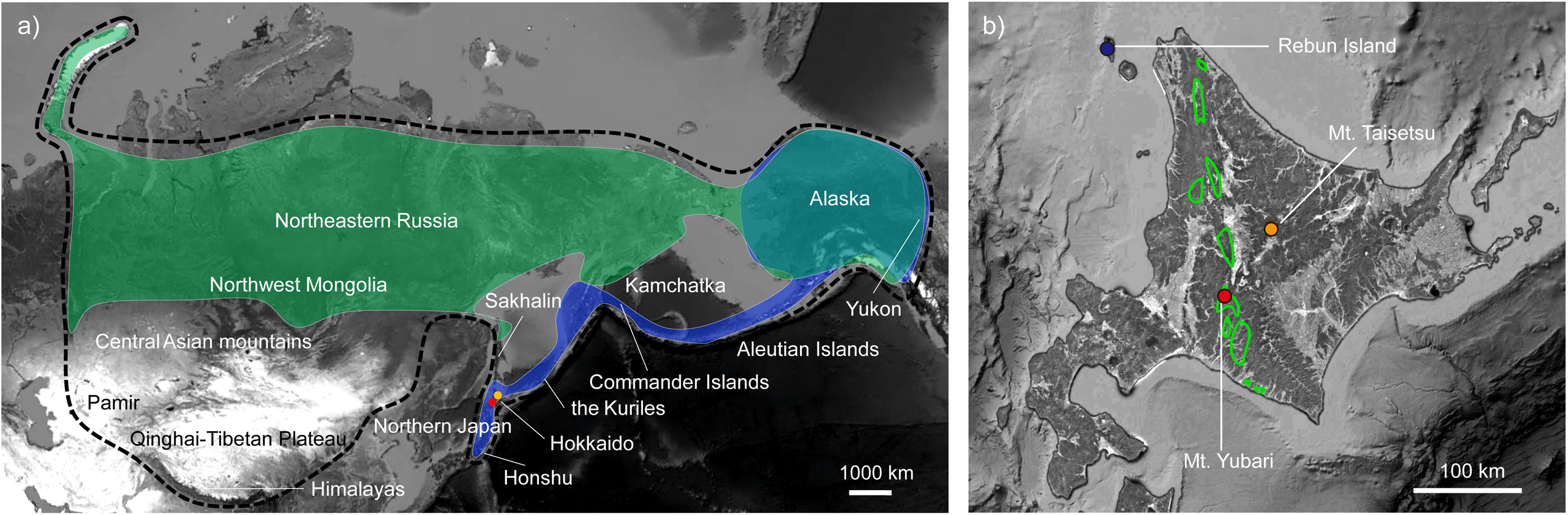
(a) Distribution areas of the genus *Lagotis* and the four-studied species, *L. takedana, L. yesoensis, L. glauca* and *L. minor*. Dotted line, the genus *Lagotis*; red, *L. takedana*; orange, *L. yesoensis*; blue, *L. glauca*; green, *L. minor*. (b) Detail distribution areas of *L. takedana, L. yeseoensis*, and *L. glauca* in Hokkaido, Japan, which are indicated by red, orange and blue, respectively. The light green areas indicate the distribution of serpentine soils of the Kamuikotan metamorphic belts.

In the present study, We aimed to resolve the phylogenetic relationships among *L. takedana, L. yesoensis, L. glauca*, and *L. minor* (as below indicated as the *Lagotis minor-glauca* complex) and to test the phylogenetic distinctiveness of the two narrow endemics, a serpentine endemics *L. takedana* and a non-serpentine endemics *L. yesoensis*. Furthermore, We reconstructed phylogeographic history of the Hokkaido endemics based on molecular dating and geological history.

## Materials and methods

### Taxon sampling

*Lagotis* J.Gaertn. is a small genus of perennial herbs comprising approximately 29 species. This genus is mainly distributed in moist sandy and gravelly areas in the Qinghai-Tibetan Plateau, Himalayas, Pamir, Central Asian mountains, arctic and subarctic Asia (northwest Mongolia, northern Japan and Russia), Alaska and the Yukon (Lu 1992; Fig. 1). In this study using cpDNA and nrITS sequence, 25 species of *Lagotis* among about 29 species were included. Four species, *L. uralensis* Schischk., *L. ikonnikovii* Schischk., *L. spectabilis* Kurz., and *L. blatteri* O.E.Schulz, were not obtained or analyzed in the present study. However, our samples included species representing all the sections and series recognized by Lu (1992) and covered the entire geographic range of *Lagotis* from northeastern Russia to Alaska.

We collected samples of *L. takedana, L. yesoensis, L. glauca* and *L. minor* from wild populations and collections in botanical gardens and herbaria. *Lagotis takedana* grows in serpentine sites with strong winds and occasional landslides at Mt. Yubari. Only two populations are known for this species, in which a few patches comprising several thousand individuals (A. Sugano personal observation). We collected samples in both the populations. *Lagotis yesoensis* grows in moist sandy and gravelly areas at Mt. Taisetsu. There are seven populations known for this species, with scattered patches containing tens of thousands of individuals, and samples were collected from five populations. *Lagotis glauca* grows in alpine areas and lowlands in the high-middle latitudes with alpine-like climates. We collected samples from two populations in Rebun Island and one population at Mt. Avacha, Kamchatka. In Rebun Island, the number of individuals of this species is only about one thousand (A. Sugano personal observation). *Lagotis minor* grows in mossy, moss-lichen, open grass tundra on slopes in the northeastern Russia to Alaska and the Yukon; this species was not collected from wild populations but from herbarium collection (*below*). In field sampling, leaves were collected from approximately twenty plants, with at least a few meters between sampled individuals, in each population for a future population genetic study and a portion of the collection was used in the present phylogenetic study. Sampling in Mt. Yubari, Mt. Taisetsu, and Rebun Island was conducted with permission from the authorities because these taxa and areas are protected by law. Living collections of Botanic Garden, Hokkaido University and Hakuba Goryu Alpine Botanical Garden were used, i.e., one samples of *L. takedana* from Mt. Yubari, three samples of *L. yesoensis* from Mt. Taisetsu, and three samples of *L. glauca* from Mt. Hakuba. Additionally, herbarium specimens were utilized to cover the species ranges. Leaf fragments were carefully collected from specimens of *L. glauca* from the Kuriles (one from each Rasshua, Chirpoi, Urup, and Onekotan islands) and *L. minor* from Schmidt Peninsula of Sakhalin in the herbarium of Hokkaido University Museum (SAPS). All the samples are described in Table 3 and Table S1. Additionally, a specimen of *L. kunawurensis* (Royle ex Benth.) Rupr. from the Himalayan range were also sampled. Sequence data for the other 20 species were downloaded from GenBank. *Wulfenia carinthiaca* Jacq. was selected as an outgroup based on a previous phylogenetic analysis of *Lagotis* (Li et al. 2014). All the other samples are listed in Table S2. In monophyletic species, that was revealed in the previous study (Li et al. 2014), sequence data were downloaded for one sample per species (in *L. integrifolia* (Wild.) Schischk., *L. integra* W.W.Sm., *L. alutacea* W.W.Sm. *L. brevituba* Maxim, *L. angustibracteata* P.C.Tsoong & H.P.Yang, *L. decumbens* Rupr., *L. globosa* Hook.f. and *L. brachystachya* Maxim.).

**Table 3.** Collection locality, population, and number of samples analyzed in cpDNA, ITS, LCN and MIG-seq of the *Lagotis minor-glauca* complex from wild populations, botanic gardens, and herbaria. A dash indicates missing data.

| Locality | Population | cpDNA | ITS | LCN | MIG-seq |
| --- | --- | --- | --- | --- | --- |
| ***L. takedana*** |  |  |  |  |  |
| Fukitoshi, Mt. Yubari, Hokkaido, Japan | Yubari1 | 3 | 3 | 3 | 12 |
| Daini Hourakuchi, Mt. Yubari, Hokkaido, Japan | Yubari2 | 3 | 3 | 3 | 12 |
| ***L. yesoensis*** |  |  |  |  |  |
| Takanegahara, Mt. Taisetsu, Hokkaido, Japan (ex-situ conservation at Botanic Garden, Hokkaido University) | Taisetsu1 | 2 | 2 | 2 | 3 |
| Mt. Aka, Mt. Taisetsu, Hokkaido, Japan (ex-situ conservation at Botanic Garden, Hokkaido University) | Taisetsu2 | 1 | 1 | 1 | 1 |
| Mt. Aka, Mt. Taisetsu, Hokkaido, Japan | Taisetsu3 | 2 | 2 | 2 | 12 |
| A branch point of the trail between Mt. Hakuun and Mt. Koizumi, Mt. Taisetsu, Hokkaido, Japan | Taisetsu4 | 2 | 2 | 2 | 10 |
| Mt. Koizumi, Mt. Taisetsu, Hokkaido, Japan | Taisetsu5 | 2 | 2 | 2 | 12 |
| ***L. glauca*** |  |  |  |  |  |
| A hill 389 m asl., Rebun Island, Hokkaido, Japan | Rebun1 | 2 | 2 | 2 | 12 |
| A hill 321 m asl., Rebun Island, Hokkaido, Japan | Rebun2 | 2 | 2 | 2 | 7 |
| Mt. Rebun, Rebun Island, Hokkaido, Japan | Rebun3 | - | - | - | 1 |
| Mt. Hakuba, Nagano, Japan (ex-situ conservation at Hakuba Goryu Alpine Plants Garden) | Hakuba1 | 1 | 1 | 1 | 2 |
| Mt. Hakuba, Nagano, Japan (ex-situ conservation at Hakuba Goryu Alpine Plants Garden) | Hakuba2 | 2 | 2 | 2 | 10 |
| Mt. Avacha, Kamchatka Peninsula, Russia | Kamcha1 | 2 | 2 | 2 | 11 |
| Rasshua Island, the Kuriles, Russia | Rasshua1 | 1 | 1 | 1 | 1 |
| Chirpoi Island, the Kuriles, Russia | Chirpoi1 | 1 | 1 | 1 | 1 |
| Urup Island, the Kuriles, Russia | Urup1 | 1 | 1 | 1 | 1 |
| The Commander Island, Kamchatka Krai, Russia | - | 1 | 1 | - | - |
| Alaska, USA | - | - | 1 | - | - |
| ***L. minor*** |  |  |  |  |  |
| Southern Schmidt Peninsula, Sakhalin, Russia | Sakhalin1 | 2 | 2 | 2 | 2 |
| Alaska, USA | - | 1 | 1 | - | - |

### DNA extraction, amplification, and sequencing

Total genomic DNA was extracted from silica-dried leaf samples using the cetyltrimethyl ammonium bromide (CTAB) method (Doyle and Doyle 1990). The quality of extracted DNA was checked by agarose gel electrophoresis. Internal transcribed spacer region of nuclear ribosomal DNA (nrITS) and four chloroplast DNA regions (*matK, rps16, trnG-trnS, trnL-F*) used in the previous phylogenetic analysis of *Lagotis* (Li et al. 2014) were hired in the present study (Table S3).

In addition, low copy nuclear (LCN) gene markers were used for selected samples that were monophyletic in the present cpDNA analyses (*Results*). We first screened 8 LCN markers that were used for other genera of Plantaginaceae, and 11 LCN markers that have been applied to a broad range of angiosperm (Table S4). Fifteen out of the 19 LCN markers did not amplify or showed multiple bands (Table S4). The other four LCN markers *LCN20, LCN38, LCN46* (Mayland-Quellhorst et al. 2016) and *Agt1* (Naumann et al. 2011) were successfully amplified and utilized (Table S5). PCR amplification was performed in 20 μl total volume: 10 μl of Taq DNA polymerase 2x master mix (Amplicon, Rødovre, Denmark), 0.8 μl of each forward and reverse primers (10 pmol/μl), 0.4 μl of DMSO, 0.8 μl of total DNA (ca. 20ng/μl) and 7.2 μl of distilled water. The PCR cycle conditions were as follows: cpDNA: initial template denaturation for 4 min at 95 °C; 25 cycles of 50 s at 94 °C, 40 s at 50 °C, and 40 s at 72 °C; a final extension for 10 min at 72 °C. nrITS: 4 min at 95 °C; 25 cycles of 50 s at 94 °C, 40 s at 60 °C, and 40 s at 72 °C; 10 min at 72 °C. *Agt1*: 2 min at 94 °C; 40 cycles of 45 s at 96 °C, 30 s at 55 °C, and 1.5 min at 72 °C; 7 min at 72 °C. *LCN20, LCN38* and *LCN46*: 2min at 98 °C; 45 cycles of 15 s at 98 °C, 30 s at 55 °C, and 30 s at 72 °C; 5 min at 72 °C. The PCR products were purified by isopropanol precipitation. The purified PCR products were amplified using ABI PRISM Big Dye Terminator v.3.1 (Applied Biosystems, Foster, CA, USA) with the same primers as those used for the PCR. Sequence reaction was performed in 20 μl total volume: 0.3 μl of BigDye Terminater 3.1 Ready Reaction Mix 2.5 X, 3.85 μl of BigDye Terminater v1.1 & v3.1 5 X Sequencing Buffer, 1μl of primer (3.2 μM), 13.85 μl of distilled water and 1 μl of PCR product. The condition of sequence reaction was as follow: initial template denaturation for 1 min at 96 °C; 25 cycles of 10 s at 96 °C, 5 s at 50 °C and 4 min at 60 °C; a final extension for 4 min at 60 °C. DNA sequencing was performed on an ABI Prism 3730 DNA analyzer (Applied Biosystems). Automatic base-calling was checked manually using Fintch TV v.1.4 (Geospiza, Seattle, WE, USA) or Chromas 2.6.2 (http://technelysium.com.au/wp/chromas/). Multiple heterozygotic sites were found in the four LCN markers (note that *L. takedana* and *L. glauca* were diploid, *L. minor* included both diploid and tetraploid, and *L. yesoensis* has not been cytologically studied; Albach et al. 2018). To statistically estimate haplotypes, PHASE 2.1 (Stephens and Donnelly 2003), as implemented in DnaSP v5 (Librado and Rozas 2009), was used with default parameter settings. A posterior probability cutoff was set to 0.7 and below this, each ambiguous site was coded as missing data. Sequence alignment was conducted using ClustalX v.2.1 (Larkin et al. 2007) and then edited manually using SeaView v. 4.6.1 (Gouy et al. 2010).

### Genome-wide single-nucleotide polymorphism genotyping (MIG-seq) analysis

We used a novel approach that was termed “multiplexed inter-simple sequence repeats (ISSR) genotyping by sequencing” (MIG-seq), which is a PCR-based procedure for single nucleotide polymorphism (SNP) genotyping using next-generation sequencing (Suyama and Matsuki 2015) to examine the phylogenetic structure the *Lagotis minor-glauca* species complex which was phylogenetically closely related based on sequence analysis (*see Results*).

The 1st PCR step was performed to amplify ISSR regions from genomic DNA with MIG-seq primer set-1 according to Suyama and Matsuki (2015).1st PCR amplification was performed in 7 μl total volume: 3.5 μl of 2x Multiplex PCR Buffer (Multiplex PCR Assay Kit Ver.2, Takara Bio, Kusatsu, Japan), 0.14 μl of each 1st PCR primers (10 μM), 0.035 ul of Multiplex PCR Enzyme Mix (Multiplex PCR Assay Kit Ver.2, Takara Bio), 1 μl of total DNA (ca. 20ng/μl) and 0.225 μl of distilled water. The 1st PCR cycle conditions were as follows: initial template denaturation for 1 min at 94 °C; 30 cycles of 30 s at 94 °C, 60 s at 38 °C, and 60 s at 72 °C; a final extension for 10 min at 72 °C. The 1st PCR products were purified with AMPure XP beads (Beckman Coulter, Pasadena, CA, USA).The 1st PCR products from each sample were diluted 41 times with deionized water and used as the template of the 2nd PCR. 2nd PCR amplification was performed in 12 μl total volume: 2.0 μl of diluted 1st PCR product, 2.4 μl of 5x PrimeSTAR GXL Buffer (Takara Bio), 0.96 μl of each dNTP, 0.24 μl of PrimeSTAR GXL DNA polymerase (Takara Bio), 1.2 μl of common forward primer and individual reverse primer (2 μM), and 4.0 μl of distilled water. The 2nd PCR cycle conditions were as follows: 12 cycles of 10 s at 98 °C, 15 s at 54 °C, and 60 s at 68 °C. The size of each 2nd PCR products (libraries) were measured and visualized using a Microchip Electrophoresis System (MultiNA, Shimadzu, Kyoto, Japan) with the DNA-2500 Reagent Kit (MultiNA). The libraries from each 2nd PCR products with those index were pooled in equimolar concentrations. The mixed libraries were purified with AMPure XP beads (Beckman Coulter) to reduce the salt concentration. Fragments of the library in the size range 300-800 bp were isolated using Pippin Prep DNA size selection system (Sage Science, Beverly, MA, USA). The final concentration was measured using a SYBR green quantitative PCR assay (library Quantification Kit; Clontech Laboratories, Mountain View, CA, USA). Finally, library was denatured using NaOH (0.2 N) and mixed with 5 % of Illumina-generated PhiX control libraries. Approximately 12 pM of the library was used for sequencing on an Illumina MiSeq Sequencer (Illumina, San Diego, CA, USA), using a MiSeq Reagent Kit v3 (150 cycle, Illumina).

### Phylogenetic analyses

To test for phylogenetic congruence between cpDNA and nrITS, the incongruence length difference test (Farris et al. 1994) was conducted using the partition homogeneity test with 100 replicates using PAUP* v.4.0a 150 (Swofford 2002). This test revealed significant heterogeneity between the data sets (*P* < 0.05), and analyses were therefore separately performed for cpDNA and nrITS data sets. Phylogenetic analyses were conducted using maximum parsimony (MP), maximum likelihood (ML), and Bayesian inference (BI).

Indels potentially have phylogenetic information. However, they often show high levels of homoplasy and can be difficult to align, especially when a wide range of taxa are sampled (Meseguer et al. 2014). Therefore, gaps were treated as missing data in this analysis. The MP analysis was performed with PAUP using heuristic searches of 1000 replicates with random taxon addition and tree bisection reconnection (TBR) branch swapping, with the MulTrees and steepest descent options on, and the MaxTrees option set to 10,000. Bootstrap support (BS; Felsenstein 1985) was estimated from 1000 replicates with random taxon addition and tree bisection reconnection branch swapping, with the MulTrees option off and the MaxTrees option set to 200. To perform ML and BI analyses, the best model of nucleotide substitution was selected with MrModeltest (Nylander 2004) using Akaike information criterion (AIC; Akaike 1974). The ML analysis was conducted with RAxML v.8.2.X. using 1000 rapid bootstrap analysis and subsequently searched for the best scoring ML tree in one program run (Stamatakis 2014). BI analysis was conducted using MrBayes 3.2.6 (Ronquist 2012). Analyses were run for 30 million generations, sampling every one thousand generations. Convergence and effective sample size (> 200 after burn-in) of all parameters were checked with Tracer v.1.6 (Drummond and Rambaut 2007) and then the first 3,000 trees were discarded as burn-in. The 50% majority rule consensus tree of all the post-burn-in trees and posterior probability (PP) was calculated. The MP, ML and BI trees were displayed using Figtree v.1.3.1 (Drummond and Rambaut 2007).

### Molecular dating analysis

To discuss historical biogeography of *Lagotis* in Hokkaido, Bayesian molecular dating was conducted based on nrITS sequence data with BEAST v.1.7.5 (Drummond et al. 2012). The best model of nucleotide substitution as selected with MrModeltest was used and base frequencies were fixed to observed frequencies in the data (i.e., an empirical base frequency setting). An uncorrelated lognormal distribution model was applied for rate variation among lineages. Speciation birth death model was employed for the branching rates. The unweighted pair-group method of arithmetic averages (UPGMA) was used to construct a starting tree. The calibration point was the split between *Lagotis* and *Wulfenia* (Mean = 19.3 Ma, 95 % highest posterior density [HPD] interval = 11.3-29.6 Ma), as estimated in Li et al. (2014), and normal-distribution age prior was used with an initial value of 19.3, mean value of 19.3, and a standard deviation of 4.1 (95 % prior interval of 11.26-27.34). Analyses were run for 30 million generations, sampling every one thousand generations. Convergence and effective sample size (> 200 after burn-in) of all parameters were checked with Tracer and then the first 3,000 trees were discarded as burn-in. A maximum clade credibility tree was estimated with a posterior probability limit of 0.5 by TreeAnnotator ver. 1.5.4 (Drummond and Rambaut 2007), and visualized with FigTree.

### Data analysis of MIG-seq

The SNP data obtained by MIG-seq were pre-processed using the FASTX-Toolkit (http://hannonlab.cshl.edu/fastx_toolkit/) enable low-quality reads removed by the ‘quality_filter’ option, using the settings of q=30 and p=40. Adapter sequence that these reads contained were removed using TagDust (Lassmann et al. 2009). To obtain SNP from the quality-filtered reads, Stacks v. 1.15 (Catchen et al. 2011) was used. The setting parameter using the ‘ustacks’ option was modified of Suyama and Matsuki (2015); maximum distance between stacks, the minimum depth of coverage required to create a stack, and maximum distance allowed to align secondary reads to primary stacks was set as 1, 5 and 1, respectively. In the ‘cstacks’ option, the parameter number of allowed mismatches between samples was set as 2.

The resulting data set were checked using the NCBI nucleotide database using blastn in BLAST + 2.2.29 (ftp://ftp.ncbi.nih.gov/blast/executables/LATEST/) with default settings and an *E*-value cutoff 10^−10^. Loci that had matched the organism that did not originated from plants were removed for further analysis.

We analyzed two different data set, for the *Lagotis minor-glauca* species complex and for the selected taxa of the *Lagotis minor-glauca* species complex that was not monophyletic based on sequence analysis (*see Results*). To exported genotype data, the ‘population’ option in Stacks was used. The exported genotype data were then processed using PLINK v.1.9 (Chang et al. 2015) and SNP loci with a minor allele frequency <0.05, missing individual rate >0.6, missing locus rate >0.1 and maximum observed heterozygosity rate >0.7 and deviation of Hardy-Weinberg equilibrium (*P* < 0.01) were filtered out.

Principal coordinate analysis (PCA) was performed to examine phylogenetic structure using GENODIVE (Meirmans and Van Tienderen 2004) with the default setting parameters.

## Results

### Phylogenetic positions of the *Lagotis minor-glauca* species complex in the genus based on cpDNA and nrITS

The aligned length of cpDNA and nrITS was 2665 and 548 bp. The numbers of parsimony informative sites of cpDNA and ITS was 48 (1.8 %) and 46 (8.4 %). In this study, clades with PP ≥0.9 and/or BS_MP_ ≥70% and/or BS_ML_ ≥70% were considered statistically supported. In either of cpDNA and nrITS markers, there was no topological incongruence among MP, ML and BI trees (data not shown). Therefore, only BI trees are presented with BS_MP_ and BS_ML_ (Fig. 2).

**Fig. 2.**
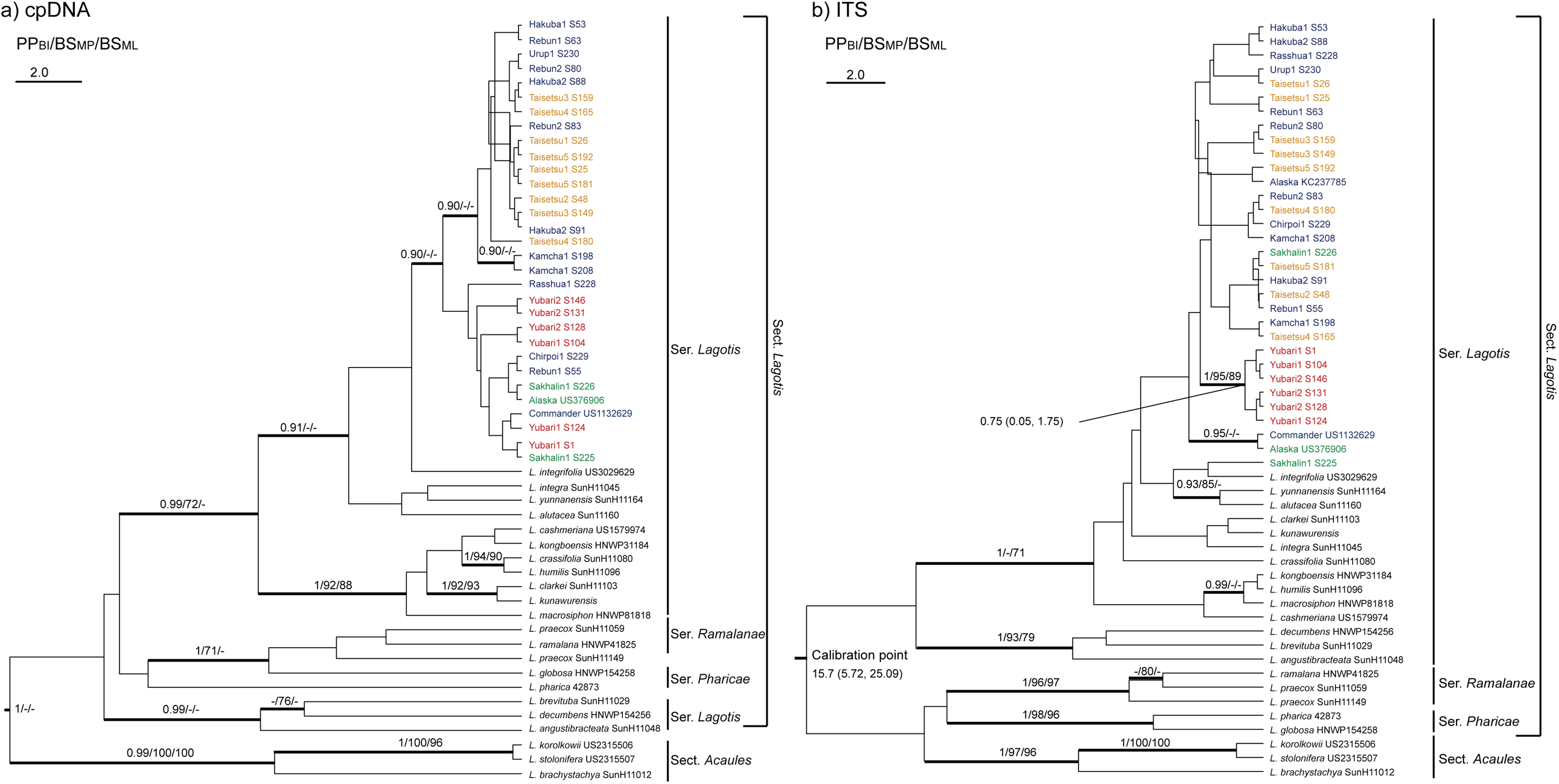
Bayesian phylogenetic tree of 25 *Lagotis* species including Hokkaido endemic *L. takedana* and *L. yesoensis* and two allied species *L. glauca* and *L. minor*. (a) and (b) are based on cpDNA and nrITS, respectively. *Lagotis takedana, L. yesoensis, L. glauca* and *L. minor* are indicated in red, orange, blue, and green, respectively. Branches in bold are supported with PP ≥ 0.9 and/or BS_MP_≥70 and/or BS_ML_≥70. Scale bar indicates 2.0 substitutions per site. For details of samples, refer Table3 and Online Resource 1.

In the cpDNA phylogenetic tree (Fig. 2a), the clade of sect. *Acaules* (PP/ BS_MP_/BS_ML_=0.99/100%/100%) was basal to the other species, that are all members of sect. *Lagotis*. Monophyly was supported neither for sect. *Lagotis* nor each of the three series *Lagotis, Ramalanae*, and *Pharicae*. Ser. *Lagotis* was divided into two clades, one including *L. takedana, L. yesoensis, L. glauca, L. minor* and other 11 species (0.99/72%/–) and the other comprising the other three species (0.99/–/–). The former clade of ser. *Lagotis* was divided into two subclades, one including *L. takedana, L. yesoensis, L. glauca*, and *L. minor* with other four species (0.91/–/–) and the other including the other seven species (1/92%/88%). Phylogenetic relationship among *L. takedana, L. yesoensis, L. glauca, L. minor*, and the four species was not resolved. *Lagotis takedana, L. yesoensis, L. glauca*, and *L. minor* formed a clade (0.90/–/–), while monophyly of each of the four species was not supported. Within the clade, *L. yesoensis* and a portion of *L. glauca* samples (two from Kamchatka, one from Urup, three from Rebun, and three from Hakuba) formed a subclade (0.90/–/–). No geographic trend was recognized in the clustering of the *L. glauca* samples.

In the nrITS phylogenetic tree (Fig. 2b), sect. *Acaules* formed a clade (1/97%/96%) but the monophyly of sect. *Lagotis* was not supported. Within sect. *Lagotis*, a clade of ser. *Ramalanae* (1/96%/97%), one of ser. *Pharicae* (1/98%/96%), and two clades of ser. *Lagotis* were recognized: one including *L. takedana, L. yesoensis, L. glauca* and *L. minor* with other 11 species (1/–/71%) and the other comprising the other three species (1/93%/79%). The relationship among *L. takedana, L. yesoensis, L. glauca, L. minor* and the 11 species was not resolved due to the low resolution of the tree. A clade of *L. takedana, L. yesoensis, L. glauca*, and *L. minor*, that was found in the cpDNA phylogeny, was not indicated. It should be noted that *L. takedana* formed a clade (1/95%/89%), while monophyly was not recovered for each *L. yesoensis, L. glauca*, and *L. minor*.

### Phylogenetic relationships of the *Lagotis minor-glauca* species complex based on LCN gene markers

The present cpDNA analyses showed that the *Lagotis minor-glauca* species complex formed a monophyletic lineage. However, the resolution of the phylogenetic relationships among these four species was very low except that *L. takedana* was monophyletic in the nrITS analyses. Hence, LCN gene markers were also used to resolve the phylogeny of the four species. *Lagotis kunawurensis* was selected as an outgroup based on cpDNA tree of this study (Fig. 2a).

The aligned length of *LCN38* and *Agt1* were 506 and 396 bp. The numbers of parsimony informative sites of *LCN38* and *Agt1* were 5 (0.99 %) and 3 (0.76 %). In each marker, there was no topological incongruence among MP, ML and BI trees (data not shown). Therefore, only ML trees are presented with BS_MP_ and PP_BI_ (Fig. 3).

**Fig. 3.**
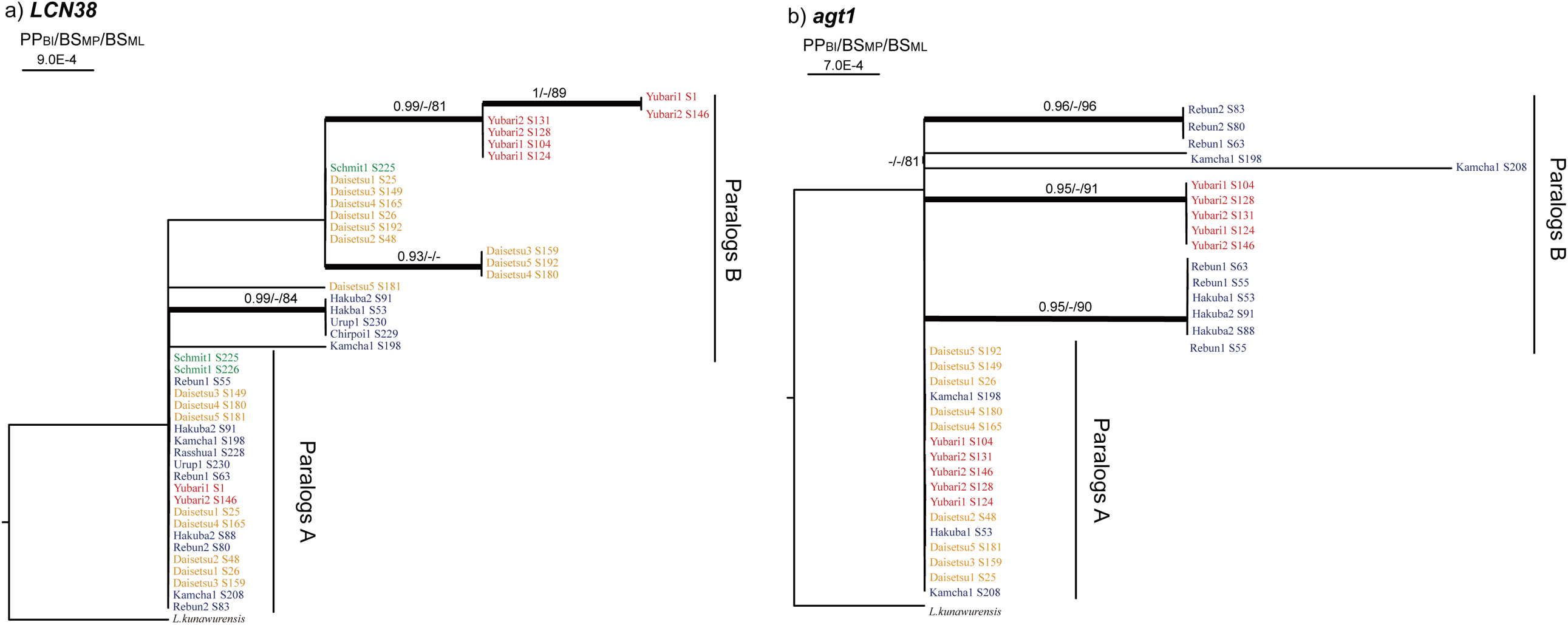
Maximum likelihood phylogenetic tree of the *Lagotis minor-glauca* species complex. (a) and (b) are based on *LCN38* and *Agt1*, respectively. *Lagotis takedana, L. yesoensis, L. glauca* and *L. minor* are indicated in red, orange, blue, and green, respectively. Branches in bold are supported with PP ≥ 0.9 and/or BS_MP_≥70 and/or BS_ML_≥70. Scale bar indicates substitutions per site. For details of samples, refer Table3 and Online Resource 1.

In the *LCN38* sequences, two paralogous copies (A and B) were recognized (Fig. 3a). It should be noted that the two paralogues were not found in all the samples; only paralogue A or B was amplified in a portion of samples of each four species, probably due to primer mismatch. In paralogue A, there was no sequence variation among the four species. On the other hand, paralogue B had comparatively large sequence variation, and monophyly of *L. takedana* was well supported, while for each of *L. yesoensis, L. glauca* and *L. minor* monophyly was not recovered. Similarly, two paralogous copies were recognized in the *Agt1* sequences (Fig. 3b). Paralogue A and B in *L. minor* and paralogue B in *L. yesoensis* were not amplified. In *L. takedana* and *L. glauca*, only paralogue A or B was amplified in a portion of samples. In paralogue A, there was no sequence variation among *L. takedana, L. glauca*, and *L. yesoensis*. In paralogue B, monophyly of *L. takedana* was supported, while monophyly was not recovered for *L. glauca*.

The aligned length of *LCN20, LCN46* were 677 and 567 bp. The numbers of parsimony informative sites of *LCN20* and *LCN46* were 3 (0.44 %) and 2 (0.35 %). In each *LCN20* and *LCN46*, multiple paralogous copies per sample were recognized. However, ML phylogenetic analysis could not determine the copy numbers of *LCN20* and *LCN46* (tree not shown). Thus, further MP and BI analyses were not conducted and these two markers were not used anymore.

### Phylogenetic structure of the *Lagotis minor-glauca* species complex based on MIG-seq

The MIG-seq analysis of the *Lagotis minor-glauca* species complex and of the selected taxa of the *Lagotis minor-glauca* species complex resulted in 552 and 519 SNPs, respectively, after filtering out the individuals and loci based on the thresholds of setting parameters as below. The variances of PCA explained by the first principle component (17.4 %) and the second one (13.9 %) were low, however, the result of PCA for the *Lagotis minor-glauca* species complex revealed *L. takedana* was clearly separated with *L. yesoensis, L. glauca* and *L. minor* (Fig. 4a), which was consistent with nrITS and two LCN phylogenetic trees. Compared with the results of cpDNA, nrITS and two LCN markers data, the phylogenetic structure of the *Lagotis minor-glauca* complex based on the MIG-seq data showed cluster of each species and geographic trends in detail. For the selected taxa of the *Lagotis glauca-minor* species complex (i.e. *L. yesoensis, L. glauca* and *L. minor*), the variances of PCA explained by the first principle component (20.0 %) and the second one (8.7 %) were low, however those revealed the genetic structure of the non-serpentine *Lagotis minor-glauca* species complex (Fig. 4b). Using the first principle component, the genetic structures revealed that *L. glauca* was separated with *L. yesoensis* and *L. minor*. Furthermore the three genetic structures was detected that *L. glauca* from the Kamchatka peninsula was located between *L. yesoensis* • *L. minor* and *L. glauca* from the Rebun lsland •Mt. Hakuba • the Kuriles. The second principle component also explained the consistency structure.

**Fig. 4.**
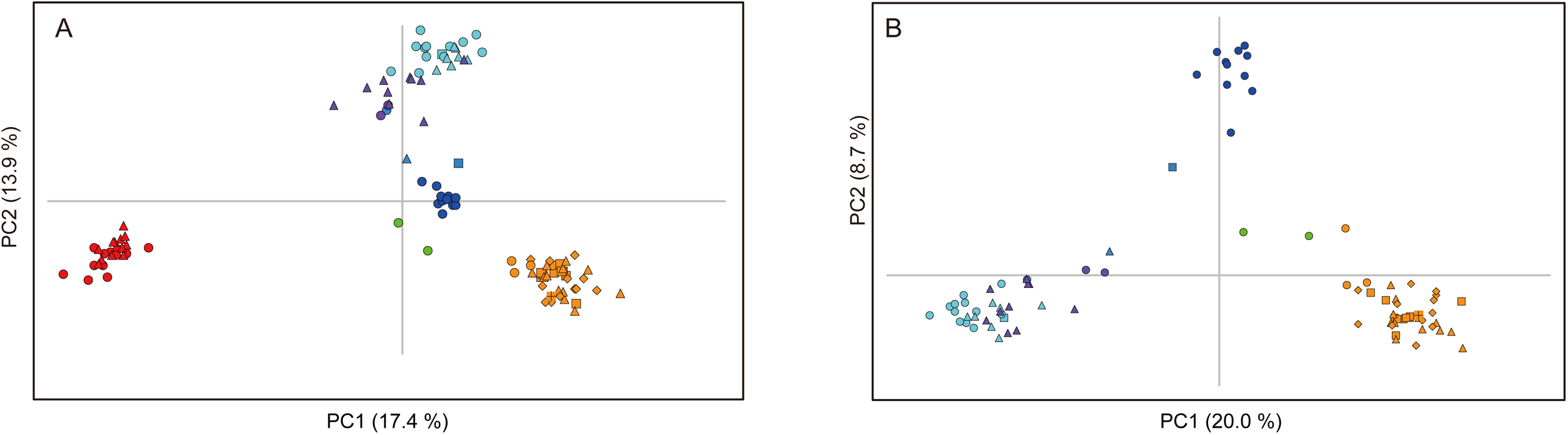
Genetic structure of the *Lagotis minor-glauca* species complex populations. Individual plots of PCA based on MIG-seq. Different populations from the same geographical localities present the same color but different markers. (a) is for the *Lagotis minor-glauca* species complex and (b) is for the species complex without *L. takedana. Lagotis takedana, L. yesoensis* and *L. minor* are indicated as red, orange and green markers, respectively. *Lagotis glauca* from Rebun Island, Mt. Hakuba, Kamchatka peninsula and the kuriles are indicated as cobalt, purple, blue and light blue markers, respectively. The percentage of total variation attributed to each axis is indicated.

### Molecular dating analysis

The Bayesian molecular dating based on nrITS estimated the age of the most recent common ancestor (MRCA) of *L. takedana* as 0.75 Ma (95 % HPD interval = 0.05-1.75 Ma). This period corresponds to the early to late Pleistocene.

## Discussion

### Phylogenetic distinctiveness of narrow endemic species, *L. takedana* and *L. yesoensis*

In the cpDNA phylogenetic tree, the *Lagotis minor-glauca* species complex formed a clade (Fig. 2a). The grouping of the *Lagotis minor-glauca* species complex reveled in this study is consistent with the morphology-based taxonomic ideas that these species are closely related (Table 2).

Monophyly of *L. takedana* was supported in the nrITS and two LCN phylogenetic trees and PCA based on MIG-seq (Figs. 2b; 3; 4a). Morphology-based taxonomy has variously treated *L. takedana* as an independent species, a variety/subspecies of *L. glauca*, or an unresolved taxon, as summarized in Table 2. The present nrITS and LCN trees and PCA of MIG-seq revealed that *L. takedana* was clearly distinguishable from *L. glauca*. Therefore, this study supports the taxonomic idea that *L. takedana* is an independent species.

In this study, *L. yesoensis* and a portion of *L. glauca* samples formed a subclade within a clade of the grouping of the *Lagotis minor-glauca* complex in the cpDNA phylogenetic tree (Fig. 2a); however, *L. yesoensis* and *L. glauca* were clearly distinguishable based on the PCA of MIG-seq (Fig. 4b). Moreover, there was no genetic differentiation between *L. yesoensis* and *L. minor* based on the PCA of MIG-seq (Fig. 4b). Our analyses of the MIG-seq data supported *L. takedana* was monophyletic, congruent with our results of nrITS and two LCN tree. However, although *L. yesoensis* and a portion of *L. glauca* samples formed a clade in the cpDNA tree, our analysese of the MIG-seq supported *L. yesoensis* and *L. glauca* was distinguishable. Given that only cpDNA data showed this incongruence and the relatively shallow of the divergent time of *L. yesoensis* and *L. glauca* (*see Discussion*), we suggested that incomplete lineage sorting may play a major role in the incongruence between cpDNA and MIG-seq data. MIG-seq is an effective method for genome-wide single-nucleotide polymorphism genotyping, thus, MIG-seq data from enough loci is possible to overcome incomplete lineage sorting owing to individual loci having different evolutionary histories. Taxonomy of *L. yesoensis* has been unclear; it has been recognized as an independent species, a subspecies/variety of *L. stelleri* (a synonym of *L. minor*), a variety of *L. minor*, or an unresolved taxon (Table 2). However, the monophyly of *L. yesoensis* was not recovered and *L. yesoensis* was not unambiguously distinguished from *L. minor* in the MIG-seq data. The result of phylogenetically indistinct *L. yesoensis* is possibly explained by (1) incorrect taxonomy, (2) hybridization with *L. minor*, or (3) incomplete linage sorting.

Concerning the first possible explanation, i.e., incorrect taxonomy, it was reported that *L. yesoensis* is morphological distinct from *L. minor* (Yamazaki 1981, 1993; Vikulova 1995; Ohashi 2017; Table 1). However, the present analyses did not separate these species. *Lagotis minor* is broadly distributed from northeastern Russia to Alaska and the Yukon. Previous morphological studies on *L. minor* examined portions of the entire species range;Vikulova (1995) chiefly investigated Russian Far East samples. Morphological comparison of *L. yesoensis* with *L. minor* should cover the entire range of the two species in order to elucidate a degree of morphological overlap. Hybridization and introgression (Rieseberg and Soltis 1991; Rieseberg et al. 1996; Ferguson and Jansen 2002; Barber et al. 2007; Pirie et al. 2009; Jabaily and Sytsma 2010; Zhang et al. 2014) and incomplete lineage sorting (Mason-Gamer et al. 1995; Wendel and Doyle 1998; Comes and Abbott 2001; Linder and Rieseberg 2004; Willyard et al. 2009) do contribute to phylogenetic incongruence (Gurushidze et al. 2010). It is sometimes difficult to distinguish between these two possible reasons in phylogenetic incongruence (Wendel and Doyle 1998; Comes and Abbott 2001; Holder et al. 2001; Jakob and Blattner 2006; Holland et al. 2008). However, the distribution area of *L. yesoensis* currently do not overlap with those of *L. minor*, and therefore the probability of hybridization is considered very low. The second possible explanation seems to be the less probable one. Given that the populations in Mt. Taisetsu (i.e. “*L. yesoensis”*) diverged from *L. minor* in a very recent time (*see below*), incomplete lineage sorting might be possible. However, our analysis based on MIG-seq data was inadequate because we used only two samples of *L. minor* and limited population. Thus, at this stage, the data are inconclusive about which explanation is more likely: incorrect taxonomy or incomplete linage sorting. Taxonomic reconsideration based on more intensive sampling, as discussed above, should be conducted. Also, further molecular analyses with rapidly evolving markers such as simple sequence repeats are needed to resolve incomplete linage sorting.

### Contrasting biogeographic history of *L. takedana* and *L. yesoensis*

The present phylogeographic analyses revealed that *L. takedana* is phylogenetically distinct while *L. yesoensis* is not. *Lagotis takedana* and *L. yesoensis* has been reported that they are endemic to Mt. Yubari and Mt. Taisetsu respectively and there are geohistorical and environmental differences between these two mountains. Mt. Yubari is located in the Kamuikotan metamorphic belt and has serpentine sites (Fig. 1b; Gouchi 1983). Mt. Yubari was formed with mountain uplift by the early Pleistocene and its alpine environment was created through it (Shimizu 1999). On the other hand, Mt. Taisetsu is volcanic and was formed through repeated volcanic activities since the Pliocene and even after the last glacial period (Kokubutani et al. 1968; Wada 2007).

In this study, *L. takedana* represents a distinct species from the others *Lagotis minor-glauca* complex. Althought the phylogenetic relationship among the *Lagotis minor-glauca* complex was not resolved cleary, the complex formed a clade, indicating that *L. takedana* is a narrow endemic species derived from either the widespread *L. glauca* or *L. minor*. Elsewise, the MRCA of the *Lagotis minor-glauca* complex led to the speciation of *L. takedana*. In the molecular dating based on nrITS, it was estimated that *L. takedana* has the MRCA in the early to late Pleistocene (mean, 0.75 Ma; 95 % confidence interval, 0.05-1.75 Ma). This period fits the time when Mt. Yubari was formed with mountain uplift and its alpine habitats became available. Mt. Yubari is one of typical examples of mountains with serpentine and limestone-derived soils (Horie 2000). In serpentine soils, high levels of endemism have been reported (Brady et al. 2005). Serpentine soil accumulates nickel and magnesium that give toxic effect to plants (Gabbrielli and Pandolfini 1984; Gabbrielli et al. 1990). Additionally, serpentine soils have low accumulation of phosphorus, nitrogen and potassium that are essential for plants (Nagy and Proctor 1997). Horie et al. (2000) reported that *L. takedana* accumulates nickel to high level and is tolerant to nickel toxicity. Therefore, *L. takedana* is adapted to serpentine soils. On the other hand, *L. glauca* is reported to have limited tolerance to serpentine soils by examining the distribution of the species in serpentine and non-serpentine sites in Mt. Hakuba (Hatano and Masuzawa 2008). Based on the present and these studies, possible scenario is that an ancestral linage being adapted to serpentine soils migrated into the alpine habitat of Mt. Yubari during the Pleistocene, and subsequently reproductively isolated from non-serpentine populations and speciated. Plants adapted to serpentine soils are called serpentine plants (Brooks 1987) and in Mt. Yubari many other endemics are serpentine plants, that are considered (but not proved with molecular phylogeny) to have been derived from non-serpentine progenitors, e.g., *Primula yuparensis* Takeda and *P. sorachiana* Miyabe et Tatew. (Primulaceae), *Anthoxanthum pluriflorum* (Koidz.) Veldkamp var. *pluriflorum* and *A. pluriflourum* var. *intermedium* (Hack.) Yonek. (Gramineae), and *Viola yubariana* Nakai and *V. brevistipulata* (Franch. et Sav.) W. Becker subsp. *brevistipulata* (Violaceae) (Sato 2007).

The age of *L. yesoensis* was not estimated because monophyly was not recovered for this species. However, the age of *L. yesoensis* seems to be very recent because the ancestral polymorphisms in the cpDNA data have been retained with a portion of *L. glauca* samples, although the age based on molecular dating analysis is unknown. In Mt. Taisetsu massive volcanic eruptions had occurred since the Early Pleistocene and even after the last glacial period (Wada 2007), suggesting that alpine plants did not migrate into and establish populations in Mt. Taisetsu until very recently. Shallow history of the alpine habitats in Mt. Taisetsu likely explains the lack of phylogenetic divergence between *L. yesoensis* and its progenitor (that is possibly *L. minor* based on the PCA of MIG-seq) due to incomplete linage sorting because incomplete linage sorting is observed at early stage of speciation (Knowles and Carstens 2007).

The migration route of *L. takedana* was not discussed because of the low resolution of the phylogenetic trees. Concerning the geographic origin of alpine plants in Hokkaido, Takahashi (2005) argued that Sakhalin and the Kuriles are two major routes via that arctic plants migrated to Hokkaido. Hidaka-Yubari alpine vegetation has connection with that of the Asian Continent via Sakhalin, while Taisetsu-Shiretoko alpine vegetation does with that of the North American Continent via the Kuriles (Tatewaki 1963, 1967). A phylogeograhic study on *Pedicularis resupinata* L. (Orobanchaceae) depicted that the same haplotypes were shared between Mt. Yubari and Sakhalin, and between Mt. Taisetsu and Kamchatka (Fujii 2003). However, the PCA of MIG-seq in this study depicted that *L. yesoensis* and *L. minor* from Sakhalin was not unambiguously distinguishable, suggesting that *L. yesoensis* was derived from the widespread *L. minor*, not from *L. glauca*. This indicates that *L. yesoensis* possibly migrated to Mt. Taisetsu via Sakhalin. To elucidate the migration route of *L. takedana* and *L. yesoensis*, population-level analyses using more nuclear gene loci or simple sequence repeat markers are needed.

Some Japanese endemic alpine plants evolved from arctic-alpine relatives that migrated southwards in the Pleistocene (e.g. Ikeda et al. 2012; DeChaine et al. 2013; Ikeda et al. 2014). However, given that the limited glaciation in East Asia during the Pleistocene (Hultén 1937), it is plausible that at least some endemics in Japan, were not recently derived from arctic-alpine plants but rather persisted in isolation for a prolonged period during the Pleistocene (Ikeda et al. 2014). Many Japanese alpine plants investigated by molecular analysis showed genetic differentiation between northern and central Japan (e.g. Senni et al. 2005; Ikeda and Setoguchi 2007; Ikeda et al. 2008; Aizawa et al. 2009; Ikeda and Setoguchi 2013), showing that populations in central Japan have persisted and experienced several cycles of cold and warm periods in the Pleistocene. However, our study showed that no genetic differentiation in *L. glauca* between Rebun Island and Mt. Hakuba, and even continuous genetic structure from central Japan to Kamchatka. Thus, an ancestral lineage of *L. glauca* in Japan may not been influenced by the Pleistocene climate change and long term isolation; their range expansion was during the recent glacial period.

In conclusion, contrasting biogeographic histories of two narrow endemics in closely related species illustrate the importance of effects of historical orogeny and ecological factors.

## Acknowledgements

We thank Seiichiro Miyamoto for assistance in field sampling and all colleagues of the Plant ecology and Systematics laboratory of Hokkaido university who provided critical discussion.The present study was supported by the Mitsui & Co.Environment Fund (R15-0067) and Grants-in-Aid for Scientific Research (16K18596).

